# Impaired Cu-Zn superoxide dismutase (SOD1) and calcineurin (Cn) interaction in ALS: A presumed consequence for TDP-43 and zinc aggregation in Tg SOD1^G93A^ rodent spinal cord tissue

**DOI:** 10.1101/220210

**Authors:** Jolene M. Kim, Elizabeth Billington, Ada Reyes, Tara Notarianni, Jessica Sage, Emre Agbas, Michael Taylor, Ian Monast, John A. Stanford, Abdulbaki Agbas

## Abstract

Impaired interactions between Calcineurin (Cn) and (Cu/Zn) superoxide dismutase (SOD1) are suspected to be responsible for the formation of hyperphosphorylated protein aggregation in amyotrophic lateral sclerosis (ALS). Serine (Ser)-enriched TDP-43 protein aggregation appears in the spinal cord of ALS animal models, and may be linked to the reduced phosphatase activity of Cn. The mutant overexpressed SOD1G93A protein does not properly bind zinc (Zn) in animal models; hence, mutant SOD1^G93A^ - Cn interaction weakens. Consequently, unstable Cn fails to dephosphorylate TDP-43 that yields hyperphosphorylated TDP-43 aggregates. Our previous studies had suggested that Cn and SOD1 interaction was necessary to keep Cn enzyme functional. We have observed low Cn level, increased Zn concentrations, and increased TDP-43 protein levels in cervical, thoracic, lumbar, and sacral regions of the spinal cord tissue homogenates. This study further supports our previous published work indicating that Cn stability depends on functional Cn-SOD1 interaction because Zn metal is crucial for maintaining the Cn stability. Less active Cn did not efficiently dephosphorylate TDP-43; hence TDP-43 aggregations appeared in the spinal cord tissue.

## INTRODUCTION

Amyotrophic Lateral Sclerosis (ALS) (a.k.a. Lou Gehrig Disease) is a non-treatable progressive neurodegenerative disease that is 100% fatal and claims about 30,000 lives per year in the USA (www.als.org). Nearly 90% of these cases are sporadic. The disease is characterized by the selective loss of motor neurons in the spinal cord, brain stem, and cerebral (motor) cortex [1]. The causative factors of the disease remain contentious at a molecular level, although the hallmark of ALS is the formation of superoxide dismutase-enriched plaques in Lewy bodies in the spinal cord [2]. Involvement of a previously discovered protein TDP-43 in ALS pathogenesis [3, 4] has recently attracted attention, resulting in several important papers published during the last decade [5-10]. Trans activating response region DNA binding protein (TDP-43)-enriched plaques are observed in ALS, and TDP-43 mutation-induced aggregates are considered as a promoting factor in ALS pathology [11]. TDP-43 phosphorylation and subsequent aggregation have been found in motor neurons [12, 13], suggesting that TDP-43 aggregation due to abnormal phosphorylation may also be relevant to reduced activity of Cn in ALS, since phosphatase Cn regulates the TDP-43 phosphorylation in both lower and higher organisms [14].

Partial inactivation of Cn in both sporadic and familial ALS patients as well as in an asymptomatic carrier of a dominant SOD1 mutation may suggest the role of Cn in the pathogenesis of ALS [15].Both Cn and SOD1 are zinc (Zn)-containing metalloproteins [16-21]. The failure of this protein-protein interaction(s) may lead to increased labile Zn accumulation, thereby causing toxicity in motor neurons [22, 23]. Therefore, involvement of Zn in the development of ALS is another critical concept to be considered since metal cations and oxidative injury have been implicated in the pathogenesis of ALS [24,25].

Elucidating these multiple factors will contribute to a basic understanding of Cn/SOD1/TDP-43 protein interactions that may occur in early stages of ALS development. This could lead to early treatment options focused on restoring normal protein-protein interactions [26].

The focus of our work is to provide some preliminary evidence that reduced Cn activity is due to weakened interactions between SOD1 and Cn in the transgenic (Tg) SOD1^G93A^ rodent (i.e., rat and mouse) model. We hypothesize that impaired bound Zn in both metallo-enzymes (i.e. SOD1 and Cn) dissociate and accumulate in cytosol, eventually inducing motor neuron death [27]. Subsequently, less functional Cn enzyme would fail to dephosphorylate TDP-43, leading to the formation of hyperphosphorylated TDP-43 aggregates that are considered to be the hallmark of neuropathology in 90% of ALS cases [12].

## MATERIALS AND METHODS

### Animals

Tg SOD1^G93A^ rat (B6.Cg-Tg(SOD1*G93A)1Gur/J) breeders were obtained from Taconic and offspring were genotyped using the protocol described in the Jackson Laboratory website https://www2.jax.org/protocolsdb/f?p=116:5:0::NO:5:P5 _MASTER_PROTOCOL_ID,P5_JRS_CODE:9877,004435). Rats were fed normal chow (Harlan Teklad rodent diet 8604 from Teklad diets, Madison, WI) and were housed in University of Kansas Medical Center (KUMC) animal facilities for six months. Animal use was approved by KUMC IACUC protocol (æ2013-2142).

### Tissues

After animals were deeply anesthetized and decapitated, their spinal cords were flushed out according to a previously published method [28]. Briefly, the spinal column was sectioned at about the 4^rd^ sacral vertebra. A 20”-22” gauge needle attached to a 10 ml syringe was inserted into the caudal opening of the spinal column. Approximately 5 ml of PBS was gently injected to flush the spinal cord from the rostral end of the severed spinal column. The cord was then sectioned into cervical, thoracic, lumbar, and sacral regions according to a template that served as a visual aid (Fig.S1). The template was constructed based on anatomic atlas of the rat spinal cord [29].

### Homogenate

All glassware was washed with strong acids (i.e., nitric acid and perchloric acid) to remove possible contamination for trace elements [30] that may interfere fluorometric zinc analysis. The spinal cord sections were weighed and placed in a pre-chilled 1 ml glass homogenizer on ice. Ten volumes of tissue homogenization buffer (0.32 M sucrose; 10mM HEPES; 0.5mM MgSO_4_; 10mM ॉamino caproic acid; pH 7.4) and the tissues (10 vol. buffer/g wet tissue) were gently homogenized on ice with 10-15 strokes (just enough to completely dissociate the tissue) using the cooled pestle. The protein concentrations were determined by the bicinchoninic acid (BCA) spectrophotometric method (Thermo Scientific, Pierce^TM^ BCA Protein Assay Kit; Catæ23227) and the homogenate aliquots were stored at -80^0^C until use.

### Western Blot Analysis

Spinal cord section homogenates were resolved in pre-cast 4-20% SDS/PAGE (Bio-Rad, mini-PROTEAN TGX^TM^ gels; Catæ456-1095) under the reducing conditions. The proteins were transferred onto a transfer membrane (Millipore, Immobilon^®^ PVDF membrane, CatæIPFL00010) and probed with anti-Calcineurin_**α**_, (Catæ C1956;Sigma Aldrich, St Louis, MO, USA; monoclonal antibody) anti-Calcineurin_**β**_, (Catæ C0581; Sigma Aldrich, St Louis, MO, USA; monoclonal antibody), anti-SOD1 (Catæ100269-1-AP; Protein Tech, USA, Rabbit polyclonal antibody), and anti-TDP-43 (Catæ 10782-2-AP; Protein Tech, USA) antibodies. Odyssey/LI-COR detection system (LI-COR Biosciences, Nebraska, USA) was used for analyzing the protein bands (Image Studio Ver.3.1).

### Zinc analysis

Spinal cord tissue homogenates were centrifuged (16K x g; 30 min at 4^0^C) in order to obtain a clear supernatant. Protein concentrations were measured by BCA assay. The supernatant samples were prepared for Zn assay analysis and the concentrations were measured by a fluorometric Zinc quantification kit (Abcam Inc., USA; Catæ ab176725) according to the manufacturer’s provided protocol. A Bio-Tek microplate reader (Model: Synergy HT, USA equipped with Gee 5 software version 2.05.5) was used for the fluorescence measurement.

### Statistical Analysis

Mann-Whitney U rank sum tests followed by the calculation of two-tailed p values were used for determining the significance between groups. SigmaPlot (ver.12.5) was used for graphing and statistical analysis.

## RESULTS

### Calcineurin protein levels were lower in Tg cervical and thoracic regions of the spinal cord

We analyzed protein levels of both the catalytic subunit of Calcineurin (Cn_A_ or PPP3CA) and the regulatory subunit (Cn_B_ or PPP3R1). We observed significantly lower catalytic subunit protein levels in the cervical (P≤0.001) and thoracic (P≤0.008) spinal cord of Tg animals, but not in the lumbar region (P≤0.130) in (Fig. 1A). There were no differences in the Cn_B-_regulatory subunit protein levels between the two groups (Fig. 1B). We were unable to harvest sufficient tissue from the sacral region to perform Cn_A_ and Cn_B_ western blot analysis.

**Fig. 1A.**
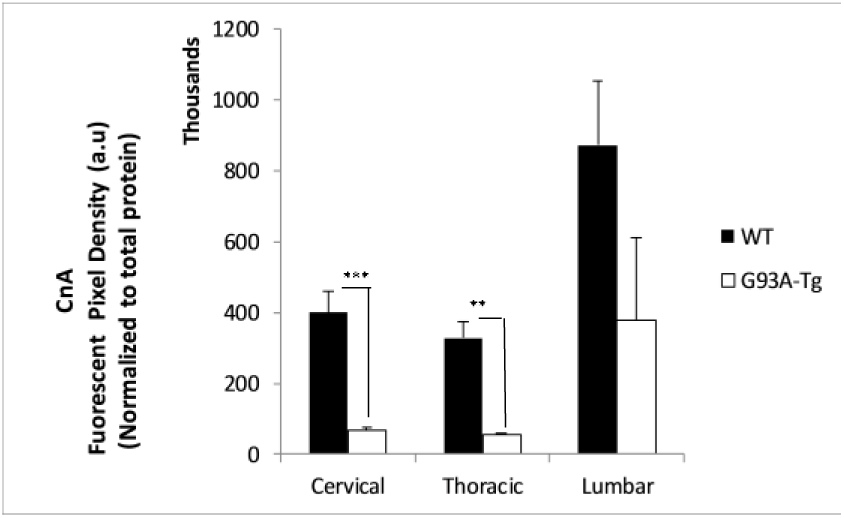
Catalytic subunit CnA protein level. Significantly lower catalytic subunit protein levi were observed in the cervical (P≤0.001) and thoracic {P≤0.OO8) spinal cord of Tg animals, bu not in the lumbar region (P≤0.130). Each bar represents the average of five animals for each group (n=5). Data are m-ean±5EM

**Fig. 1B.**
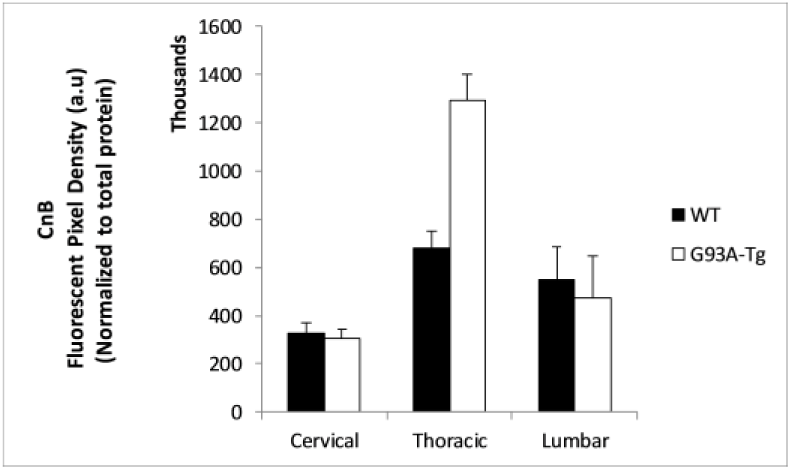
Regulatory subunit CnB protein levels. Mn differences were observed In the Cnt regulatory subunit protein levels between the two groups. Data are mean ±SEM (n-5)

### TDP-43 protein levels in spinal cord regions

We measured spinal cord TDP-43 protein levels by Western blot analysis. We observed an increasing trends of TDP-43 in thoracic, lumbar, and sacral regions; however, no statistically significant differences in TDP-43 between Tg and WT rats in any of the regions measured (Fig-2).

**Fig. 2.**
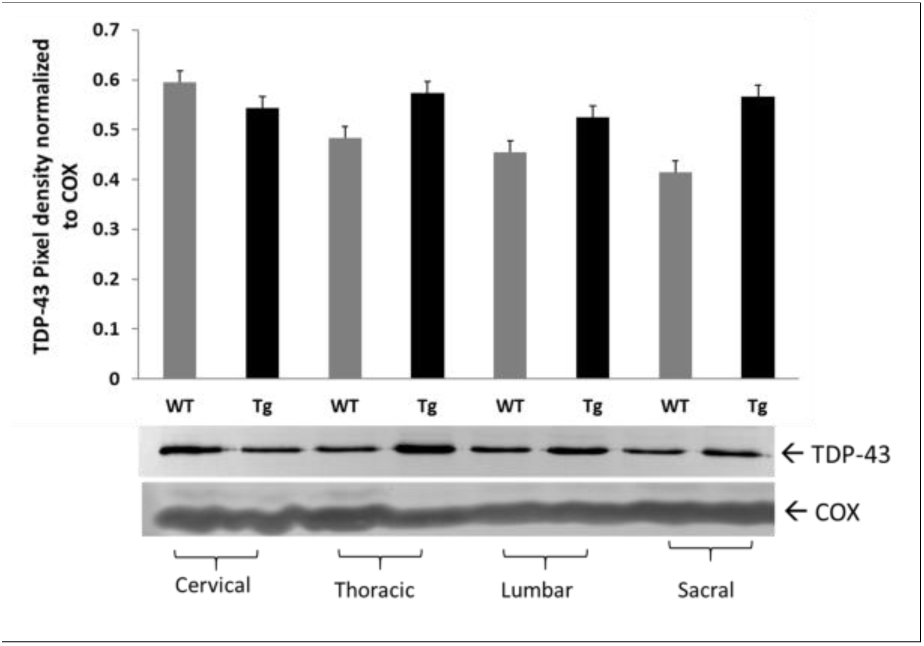
TDP-43 protein levels in spinal cord regions of SOD1MiA Tg rat. TDP43 protein levels in spinal cord regions of WT (n=6) and Tg (n=8) animals. Pan anti-TDP-43 Ab was used for Western blot analysis. No statistically significant difference was observed in cervical (P<0.0663), in thoracic (P<0.250), in lumbar (P<0.393), and in sacral (P<0.157) regions of WT and Tg animals.Data are meant SEM

### Predicted phosphorylation site analysis for TDP-43

We have employed a computer based **P**redictor **o**f **N**atural **D**isordered **R**egion (PONDR^®^) algorithm using TDP-43 sequence (NCBI accession code: Q5R5W2.1). Disordered Enhanced Phosphorylation Predictor (DEPP) analysis predicted 28 potential phosphorylation sites and a majority of them were Ser amino acid enriched on the C-terminus (aa 369-410) (Fig.S5).

### Higher Zn concentrations were measured in Tg lumbar spinal cord

We measured the total Zn concentrations in all regions of rat spinal cord (Fig-3). We measured significantly greater Zn in the lumbar region of Tg animals (P≤ 0.01). The greater Zn levels in the cervical, thoracic, and sacral regions did not reach statistical significance.

**Fig. 3.**
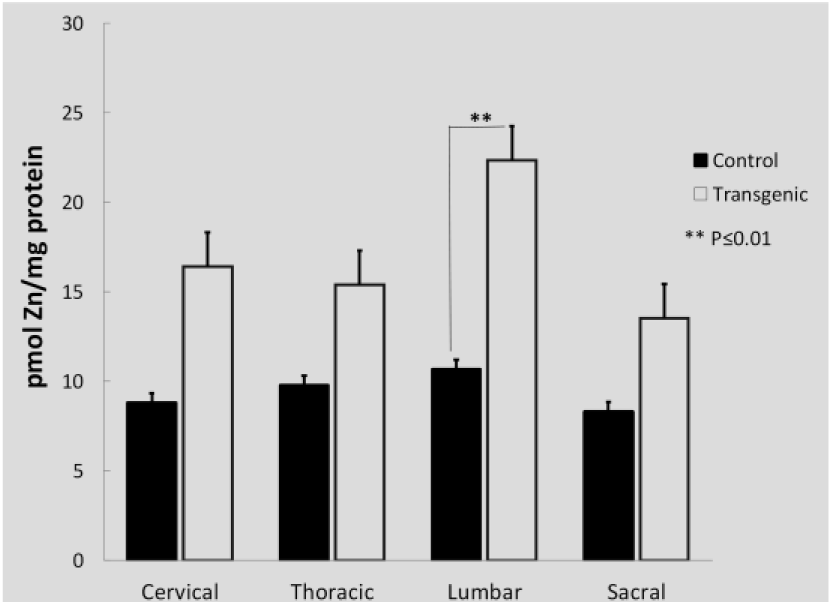
Zinc concentrations in rat spinal cord regions. Significantly greater Zn concentrations were measured In the lumbar region of Tg animals (** P≤0.01). The Zn levels In the cervical, thoracic, and sacral regions did not reach statistical significance. Data are mean±SEM (n=5)

## DISCUSSION

We previously showed that Cn activity depends on the presence of normal homodimer SOD1 under in vitro conditions and concluded that Cn-SOD1 interactions actually protect SOD1 dimers from oxidative modification [31]. Evidence provided in the literature suggests that depletion of Zn likely leads the SOD1 protein to undergo aggregation under oxidative conditions [32, 33] We have hypothesized that (i) the mutation ins SOD1^G93A^ weakens Zn binding in SOD1 [34]; leading the protein misfolding;(ii) this abnormal SOD1 may not efficiently interact with Cn, promoting Zn loss and reduced activity in Cn [31]; (iii) less active Cn fails to dephosphorylate TDP-43, leading TDP-43 hyperphosphorylation; (iv) failed SOD1-Cn interactions contributes to Zn accumulation.Data provided in this study are in agreement with this hypothesis. We have observed reduced Cn protein amounts in rat spinal cord samples as determined by Western blot analysis (Fig.1A, 1B). In unpublished data, we also observed reduced Cn phosphatase activity levels in Tg SOD1^G93A^mice (Fig.S2). We did not analyze Cn enzyme activity levels in spinal cord homogenates in rat due to insufficient amount of samples; however, we predict that low Cn levels reflect low Cn ativity. We have tested both Cn_A_ (∼61 kDa calmodulin-binding catalytic subunit) and Cn_B_ (∼19 kDa Ca^2+^/Calmodulin-binding regulatory subunit) protein levels in spinal cord regions and found that Cn_A_ protein level was significantly lower in Tg SOD1^G93A^ rat spinal cord regions as compare to WT (Fig.1A). On the other hand, Cn_B_ subunit levels did not differ between the two groups (Fig.1B).Although we observed a Cn_B_ upregulation pattern in thoracic region of Tg animals, this increase did not reach statistically significant level. Cn_B_ subunit is involved in the apoptotic reaction cascade in neurodegeneration [34]. We expected that Cn_B_ subunit upregulation would be observed spinal cord. Based on these observations, we propose that Cn-SOD1 interaction fails due to alterations in the SOD1^G93A^ mutation, then the less active Cn fails to dephosphorylate TDP-43; consequently, hyperphosphorylated TDP-43 begins to aggregate (Fig.4). Because the impaired SOD1-Cn interaction is not stabilized, Zn ions would be dissociated from both Cn and SOD1 and form aggregates. The molar ratio between SOD1:Cn was determined to be 2:1 in in vitro conditions for proper Cn activity [31].

**Fig. 4.**
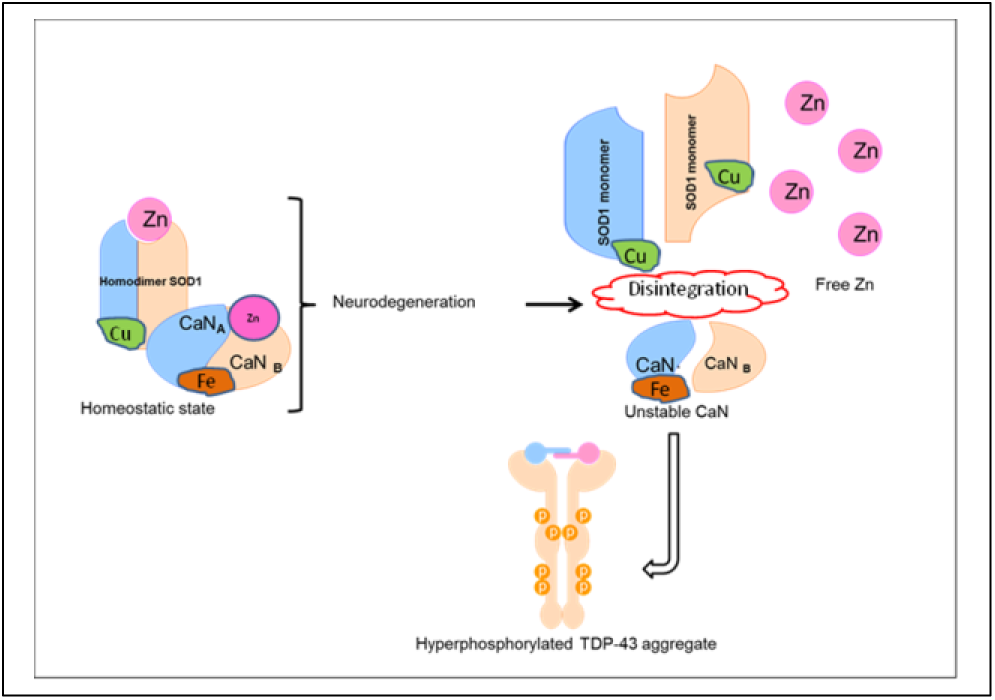
Proposed model for Cn-SOD1 impaired interaction that leads for TDP-43 aggregation and Zn accumulation in ALS

Even though TgSOD1^G93A^ rat expresses more SOD1 (Fig.S3), Cn levels remained low in thoracic and lumbar spinal cord regions. Decreased Cn enzyme activity in lymphocytes from ALS patients was also reported in the literature [15].

Recent studies have suggested that Zn dyshomeostasis occurs in spinal cords of SOD1^G93A^ transgenic mouse and may contribute to motoneuron degeneration [35, 36]; however, these studies do not specify the region(s) of the spinal cord where Zn accumulation is prevalent. In this study, we found that greater Zn concentration is present in the lumbar spinal cord of SOD1^G93A^ rats (Fig.3) and mice (Fig.S4).

The labile Zn aggregation in neurons and astrocytes in the spinal cord of SOD1^G93A^ transgenic mice supports the concept of the Zn dyshomeostasis in ALS [22]. Zinc is an essential cofactor for enzymatic activity and 10% of all mammalian proteins require Zn for proper folding [37]. Although Zn modulates the function of many other proteins, its homeostasis is poorly understood in health and in disease conditions. The elevated toxic levels of trace element Zn were reported in ALS patients reviewed by Smith and Lee [38], suggesting the link between environmental causes and an increased incidence of the disease. A possible approach would be to study the metal binding properties of SOD1, since SOD1 is a metalloprotein and binds to both Cu and Zn. The G93A SOD1 mutation selectively destabilizes the remote metal binding region (i.e., Zn and Cu binding region) [39]. We have proposed a model (Fig.4) where unstable Zn–deficient SOD1 proteins may destabilize Cn that weakly binds Zn [31], causing impaired SOD1-Cn interaction. Subsequently, destabilized Cn fails to dephosphorylate TDP-43, leading to TDP-43 protein aggregation. However, this model needs to be tested in cell-based studies. Alterations in metal-binding sites of another mutant SOD1^G85R^ contributes to toxicity in SOD1-linked ALS [40]. This observation further supports the notion that Zn deficiency may contribute to the less stable SOD1 which enhances SOD1 aggregation in neuronal cells. This hypothesis was recently challenged by reports that SOD1 stabilization is not due to dimerization and proper metallation but disulfide formation between two SOD1 homodimer [40, 41]. In addition, liberated Zn may create local metal toxicity. There are extensive reports identifying the neurotoxic effects of Zn, however how Zn kills neurons remain unclear [42].

Calcineurin is a Ser/Thr specific phosphatase [16]. Ser amino acid enriched C-terminus region of TDP-43 (Fig.S5) is a potential substrate for Cn. Therefore, decreased activity of Cn may impair the dephosphorylation of TDP-43; hence, TDP-43 aggregations form in SOD1^G93A^ Tg rat spinal cord tissue (Fig.2; Fig.S6).

In conclusion, the complex biology of ALS necessitates the study of relevant biochemical events so that logical connections among the biomolecules and metal ions can be established. In this study, we have demonstrated that impaired SOD1^G93A^ – Cn interaction gave rise to three important consequences in vivo; (1) reduced Cn enzyme levels, (2) labile Zn accumulation, and (3) TDP-43 aggregation. These events would potentially have contributed to ALS progression in the rat model. Further studies should design effective small molecules that maintain Cn activity so that hyperphosphorylated TDP-43 aggregation is controlled if not stopped. Stabilized Cn would not contribute labile Zn after dissociation from Cn molecule; therefore, Zn aggregation would be relatively less.

## ACKNOWLEDGEMENTS

We thank Justine Thomas, Megan Liberty, Tess Lamack, Mina Tawadrous, and Prabhal Sandhu for their contributions in experimental phase of this manuscript. Financial support is from KCU intramural grants awarded for AA and from GM103418 to JAS. Authors declare no financial conflicts.

